# Reproducible Brain Entropy (BEN) Alterations During Rumination

**DOI:** 10.1101/2024.12.02.626334

**Authors:** Jue Lu, Donghui Song, Da Chang, Ze Wang

## Abstract

Rumination, characterized by recurrent and repetitive thinking, is closely associated with mental disorders such as depression. However, the neural mechanisms underlying this mental state remain poorly understood. In this study, we use a relatively novel neuroimaging analysis method-Brain Entropy (BEN) to quantitatively assess the irregularity, disorder, and complexity of brain activity, providing new insights into the neural mechanisms of rumination.

We utilized a publicly available MRI dataset from three different scanners. The dataset included 41 healthy adult participants who completed identical fMRI tasks on IPCASGE, PKUGE, and PKUSIEMENS scanners. The time interval between the two visits was 22.0 ± 14.6 days. The fMRI session included four runs: resting state, sad memory, rumination, and distraction. Whole brain voxel-wise BEN differences of task state and resting state, rumination and sad memory, distraction and sad memory, and rumination and distraction were tested and overlap regions after thresholded (p<0.05) across the three scanners were identified as exhibiting significant differences.

The results demonstrate distinct alterations in BEN across mental states. Compared to the sad memory condition, decreased BEN was found in the visual cortex (VC) during rumination and decreased BEN in the posterior cingulate cortex/precuneus (PCC/PCu) during distraction. However, when compared to distraction, rumination showed increased BEN in the PCC/PCu. These findings suggest that rumination involves heightened internal focus and reduced processing of external environmental information. This study highlights BEN as a valuable metric for elucidating the neural mechanisms underlying rumination and its role in depression.

## 1. Introduction

Rumination is termed as recurrent and repetitive thinking on symptoms, feelings, problems, upsetting events, and negative aspects of the self, typically with a focus on their causes, circumstances, meanings, and implications (Watkins 2023). Rumination is closely related to depression and considered as a common mechanism relating depressive risk factors to depression (Spasojević and Alloy 2001, Whisman, du Pont et al. 2020). Importantly, rumination is not only related to depression, but is involved in the development and maintenance of a broad range of disorders, including postLtraumatic stress disorder (PTSD), anxiety disorders, insomnia, eating disorders, somatic symptom disorder, and substance use disorders and is increasingly recognized as a transdiagnostic phenomenon (Nolen-Hoeksema, Wisco et al. 2008, McLaughlin and Nolen-Hoeksema 2011, Ehring 2021, Mendoza, Mordeno et al. 2022). A recent review (Watkins and Roberts 2020) shows that rumination has multiple negative consequences: 1) exacerbating psychopathology by magnifying and prolonging negative mood states, interfering with problem-solving and instrumental behavior and reducing sensitivity to changing contingencies; 2) acting as a transdiagnostic mental health vulnerability impacting anxiety, depression, psychosis, insomnia, and impulsive behaviors; 3) interfering with therapy and limiting the efficacy of psychological interventions; 4) exacerbating and maintaining physiological stress responses. Understanding the neural substrates of rumination will greatly increase our understanding of psychopathology and facilitate subsequent treatment. However, little is known about neural substrates of rumination. The development of non-invasive neuroimaging technology provides the possibility to explore the neural substrates of rumination, but few neuroimaging studies (Denson, Pedersen et al. 2009, Burkhouse, Jacobs et al. 2017, Steinfurth, Alius et al. 2017, Apazoglou, Küng et al. 2019, Nejad, Rotgé et al. 2019) have explored rumination compared to other cognitive areas, such as attention regulation or memory, relatively (Zhou, Chen et al. 2020). Recently, a meta-analysis shown that the default mode network (DMN) may be the key brain network underlying rumination (Zhou, Chen et al. 2020) and then a recent study has revealed stable within-DMN functional connectivity (FC) was generally decreased during rumination as compared to the distraction state (Chen, Chen et al. 2020). However, it remains unclear how ruminations alter regional brain dynamics activity. This study aimed to assess the neural substrates of rumination identified by regional brain dynamics activity by fMRI-derived regional brain entropy (BEN) (Wang, Li et al. 2014) using public access dataset from RMP Rumination fMRI Dataset (http://rfmri.org/RuminationfMRIData).

The brain is considered as complexity dynamic system, entropy indicates irregularity, disorder and complexity of a dynamic system, BEN quantitative underlying irregularity, disorder, uncertainty, and complexity of regional brain activity (Wang, Li et al. 2014). The fMRI-based BEN is associated with neurocognition (Song, Lin et al., Wang 2021, Del Mauro and Wang 2024) and task performance (Lin, Chang et al. 2022, Song and Wang 2024) and is sensitive to neurochemical signal (Song and Wang 2024, Song and Wang 2024), neuromodulations (Chang, Zhang et al. 2018, Song, Chang et al. 2019, Liu, Song et al. 2024, Song, Deng et al. 2024), pharmacological (Chang, Song et al. 2018, Liu, Song et al. 2020) and non-pharmacological intervention (Dong-Hui Song 2024).The BEN also has been shown to be independent of regional perfusion and the amplitude of low frequency fluctuations of resting state fMRI in most parts of the brain cortex (Song, Chang et al. 2019), suggesting that BEN may be a special property of brain activity. More importantly, disease-related changes in BEN have been widely observed in psychiatry and neurological disorders (Li, Fang et al. 2016, Zhou, Zhuang et al. 2016, Lin, Lee et al. 2019, Xue, Yu et al. 2019, Liu, Song et al. 2020, Wang and Initiative 2020, Fu, Liang et al. 2023, Jiang, Cai et al. 2023, Liu, Ge et al. 2023, Dong-Hui Song 2024, Lin, Lee et al. 2024), including depression (Lin, Lee et al. 2019, Liu, Song et al. 2020, Dong-Hui Song 2024, Lin, Lee et al. 2024). In depression, we found that more severe depressive symptoms were associated with higher BEN in the medial orbital-frontal cortex (MOFC), which was suppressed by pharmacological treatment in patients with major depressive disorder (MDD) (Liu, Song et al. 2020). The higher BEN was observed in the prefrontal cortex in mild to moderate depression, which decreased following non-pharmacological interventions (Dong-Hui Song 2024). In healthy individuals, we found that high-frequency rTMS targeting the left DLPFC-commonly used as a protocol for treating MDD—could reduce BEN in the MOFC (Song, Chang et al. 2019). Based on that FC is negatively correlated with BEN (Song 2024) and the well-established link between rumination and DMN, along with findings from the same dataset indicating stability within-DMN FC was generally decreased during rumination, we hypothesize that BEN may increase in DMN during rumination. The RMP Rumination fMRI Dataset, collected using identical experimental protocols across three different MR scanners from GE and Siemens to enhances the reproducibility of findings—a longstanding focus in neuroimaging research (Esteban, Markiewicz et al. 2019, Marek, Tervo-Clemmens et al. 2022, Niso, Botvinik-Nezer et al. 2022, Botvinik-Nezer and Wager 2023). In this study, we investigated the neural substrates of rumination using BEN, emphasizing the reproducibility of findings obtained from data acquired using different MRI scanners.

## 2. Methods

### 2.1 Dataset

This study all data from RMP Rumination fMRI Dataset (http://rfmri.org/RuminationfMRIData). This dataset including 41 young healthy adult participants (22 females; mean age = 22.7 ± 4.1 years), each participant was scanned on 3 different MRI scanners from Chinese Academy of Sciences (IPCAS) and Peking University (PKU) under resting state, sad memory, rumination state and distraction state.

#### 2.1.1 Experimental design

Identical fMRI tasks were completed by all participants on 3 different MRI scanners with an order of counter-balanced across participants. Interval time between 2 sequential visits were 22.0 ± 14.6 days. The fMRI session included 4 runs: resting state, sad memory, rumination state and distraction state. Sad memory from 4 individual negative autobiographical events. The rumination state was introduced as “passive and repetitive thinking on negative events and their possible consequences,” and distraction state was introduced as “imagining images unrelated to yourself.” An 8-minute resting state as a baseline first came. All mental states (sad memory, rumination, and distraction) contained four randomly sequentially presented stimuli except for the resting state. Each stimulus lasted for 2 min, and then was switched to the next without any inter-stimuli intervals (ISI), forming an 8-minute continuous mental state. The resting state and negative autobiographical events recall were sequenced first and second while the order of rumination and distraction states was counter-balanced across participants.

#### 2.1.2 MRI data acquisition

Images were acquired from three different scanners. 3 Tesla GE MR750 scanners at the Magnetic Resonance Imaging Research Center, Institute of Psychology, Chinese Academy of Sciences (IPCASGE) and Peking University (PKUGE) with 8-channel head-coils. Another 3 Tesla SIEMENS PRISMA scanner (PKUSIEMENS) with an 8-channel head-coil in Peking University was also used. Scan parameters followed the recommended standardized sequence of the Association of Brain Imaging (www.abimaging.org) and these parameters were developed to harmonize site effects from different models of scanners.

All participants underwent a 3D T1-weighted scan first before functional image acquisitions with flowing parameters: 192 sagittal slices, TR = 6.7 ms, TE = 2.90 ms, slice thickness/gap = 1/0mm, in-plane resolution = 256 × 256, inversion time (IT) = 450ms, FOV = 256 × 256 mm, flip angle = 7°, average = 1 from IPCASGE and PKUGE; 192 sagittal slices, TR = 2530 ms, TE = 2.98 ms, slice thickness/gap = 1/0 mm, in-plane resolution = 256 × 224, inversion time (TI) = 1100 ms, FOV = 256 × 224 mm, flip angle = 7°, average=1 from PKUSIEMENS. Then, functional images were obtained for the resting state and all three mental states (sad memory, rumination and distraction) with flowing parameters: 33 axial slices, TR = 2000 ms, TE = 30 ms, FA = 90°, thickness/gap = 3.5/0.6 mm, FOV = 220 × 220 mm, matrix = 64 × 64 from IPCASGE/PKUGE; 62 axial slices, TR = 2000 ms, TE = 30 ms, FA = 90°, thickness = 2 mm, multiband factor = 2, FOV = 224 × 224 mm from PKUSIEMENS.

For more detailed experimental design and data acquisition information, please refer to the original article (Chen, Chen et al. 2020) and the dataset (http://rfmri.org/RuminationfMRIData).

### 2.2 MRI preprocessing and BEN mapping

#### 2.2.1 MRI preprocessing

MR images preprocessing was performed using Statistical Parametric Mapping (SPM12, WELLCOME TRUST CENTRE FOR NEUROIMAGING, London, UK, http://www.fil.ion.ucl.ac.uk/spm/software/spm12/) (Friston, Holmes et al. 1994). The following preprocessing was performed: 1) The first 4 volumes were discarded to allow the signal to reach a steady state; 2) The remaining images were performed slice timing correction; 3) Realignments on functional images were performed using a six-parameter (rigid body) linear transformation to correct head motions; 4) Structural images were segmented into grey matter (GM), white matter (WM),and cerebrospinal fluid (CSF), functional images were spatially co-registered with structural images and WM and CSF segmentation maps were back-registered into the functional images space and resampled to same resolution for extracting the mean WM and CSF signals; 5) Temporal nuisance correction was performed by regressing out six head motion parameters, WM signal, and CSF signal, global signal regression wasn’t performed; 6) Temporal bandpass filtering (0.01–0.1 Hz) was performed; 7) The functional images were smoothed with an isotropic Gaussian kernel with a full-width-at -half-maximum (FWHM) of 6 mm.

#### 2.2.2 BEN mapping

BEN mapping was performed using the BENtbx (https://cfn.upenn.edu/zewang/software.html) (Wang, Li et al. 2014). The BEN maps were calculated from the preprocessed functional images at each voxel using sample entropy (Lake, Richman et al. 2002). In this study, we follow our previous study parameter settings (Wang, Li et al. 2014, Zhou, Zhuang et al. 2016, Song, Chang et al. 2019, Song, Chang et al. 2019, Xue, Yu et al. 2019, Liu, Song et al. 2020, Wang and Initiative 2020, Wang 2021, Lin, Chang et al. 2022, Jiang, Cai et al. 2023, Song Donghui 2023, Del Mauro, Sevel et al. 2024, Dong-Hui Song 2024, Liu, Song et al. 2024, Song, Deng et al. 2024, Song and Wang 2024, Song and Wang 2024), setting the window length to 3 and the cutoff threshold to 0.6. More details of BEN calculation can be found in the original BENtbx paper (Wang, Li et al. 2014). The BEN maps were normalized to the Montreal Neurological Institution (MNI) standard space and resampled with a resolution of 3 × 3 × 3 mm^3^ and then smoothed with an isotropic Gaussian kernel (FWHM = 10 mm^3^).

### 2.3 Statistical analysis

We first estimated the differences between task-state BEN and resting-state BEN using paired t-tests, applying an initial threshold of p < 0.05. Regions overlapping across the three scans were deemed to exhibit significant differences between task-state BEN and resting-state BEN. Subsequent analyses were restricted to brain regions showing significant differences between task-state BEN and resting-state BEN. The contrasts of rumination vs. sad memory, distraction vs. sad memory, and rumination vs. distraction were evaluated using paired t-tests for each scan. An initial threshold of p < 0.05 was set, and regions consistently overlapping across the three scanners were identified as exhibiting significant differences. Finally, the mean BEN values for regions with significant differences were extracted, and individual-level variability was visualized through plotting and were tested using paired t-tests. These analyses were conducted using customized Python scripts based on Nilearn (https://nilearn.github.io/stable/index.html).

## 3. Results

Compared to the resting state, task states exhibit higher BEN in the dorsolateral prefrontal cortex (DLPFC), temporo-parietal junction (TPJ), posterior cingulate cortex and precuneus (PCC/PCu), along with lower BEN in the visual cortex (VC) (Fig.1A). Compared to sad memory, lower BEN in the visual cortex (VC) (Fig.1B) during ruminations, while lower BEN in the PCC/PCu during distractions (Fig.1C). Additionally, higher BEN during ruminations in the PCC/PCu compared to distraction (Fig.1D). Subsequently, the mean BEN values were extracted from the VC and PCC/PCu for each subject during each scan. The results showed significant differences across the three-task state from three scanners, with BEN during rumination being significantly lower than during sad memory in the VC. Compared to distraction, rumination exhibited higher BEN in the PCC/PCu, while distraction showed lower BEN in the PCC/PCu relative to sad memory (Fig.2).

**Fig. 1.**
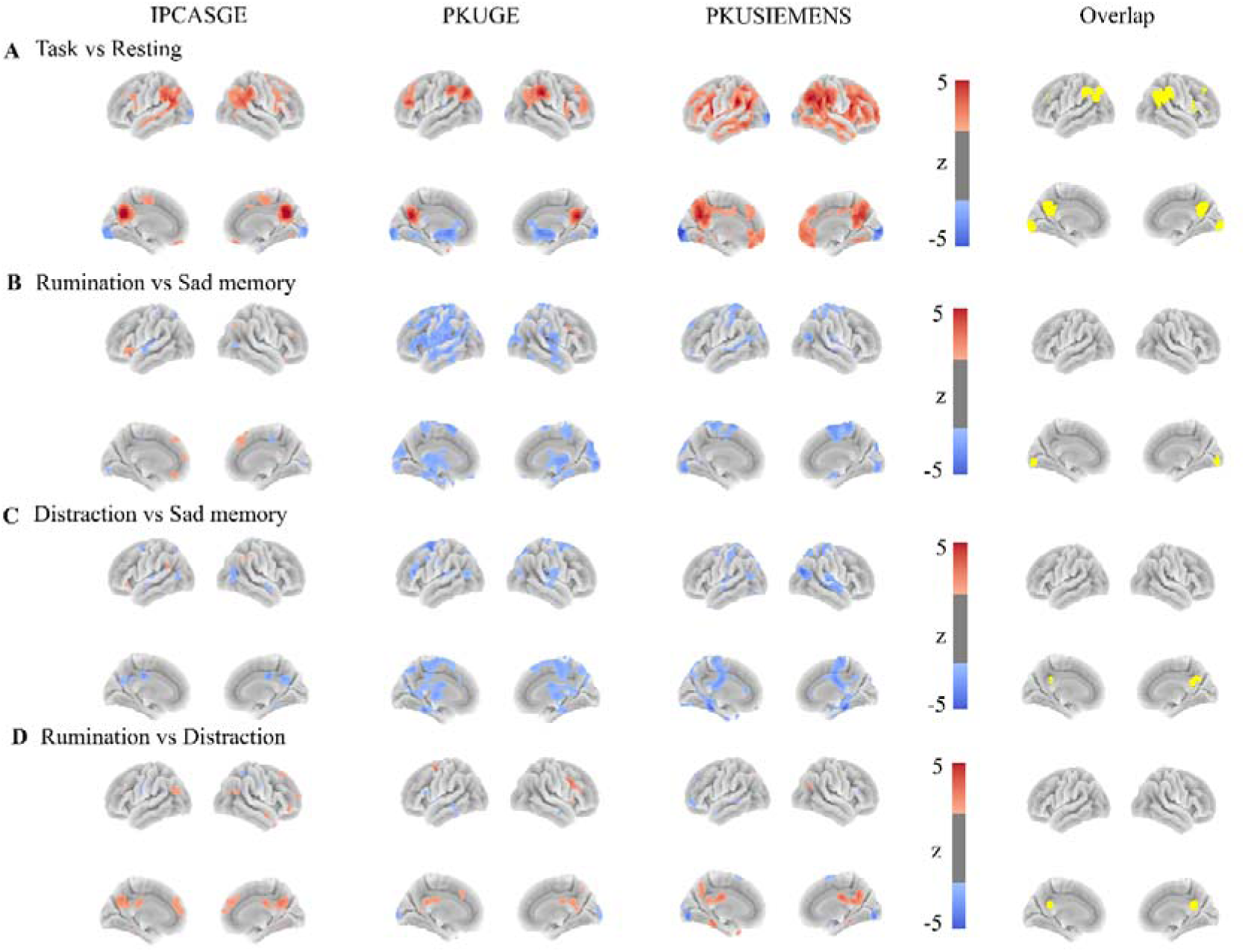
The differences in BEN between different task states. A. The differences between task-state BEN and resting-state BEN were analyzed separately for IPCASGE, PKUGE, and PKUSIEMENS, with the overlapping brain regions across the three scans identified and highlighted. The colorbar indicates z score, warm color means higher BEN during task-state, cool color means lower BEN during task-state. B. The differences in BEN between the rumination and the sad memory were analyzed separately for IPCASGE, PKUGE, and PKUSIEMENS, with the overlapping brain regions across the three scans identified and highlighted. The colorbar indicates z score, warm color means higher BEN during ruminations, cool color means lower BEN during ruminations. C. The differences in BEN between the distraction and the sad memory were analyzed separately for IPCASGE, PKUGE, and PKUSIEMENS, with the overlapping brain regions across the three scans identified and highlighted. The colorbar indicates z score, warm color means higher BEN during distractions, cool color means lower BEN during distractions. D. The differences in BEN between the rumination and distraction were analyzed separately for IPCASGE, PKUGE, and PKUSIEMENS, with the overlapping brain regions across the three scans identified and highlighted. The colorbar indicates z score, warm color means higher BEN during ruminations, cool color means lower BEN during ruminations.

**Fig. 2.**
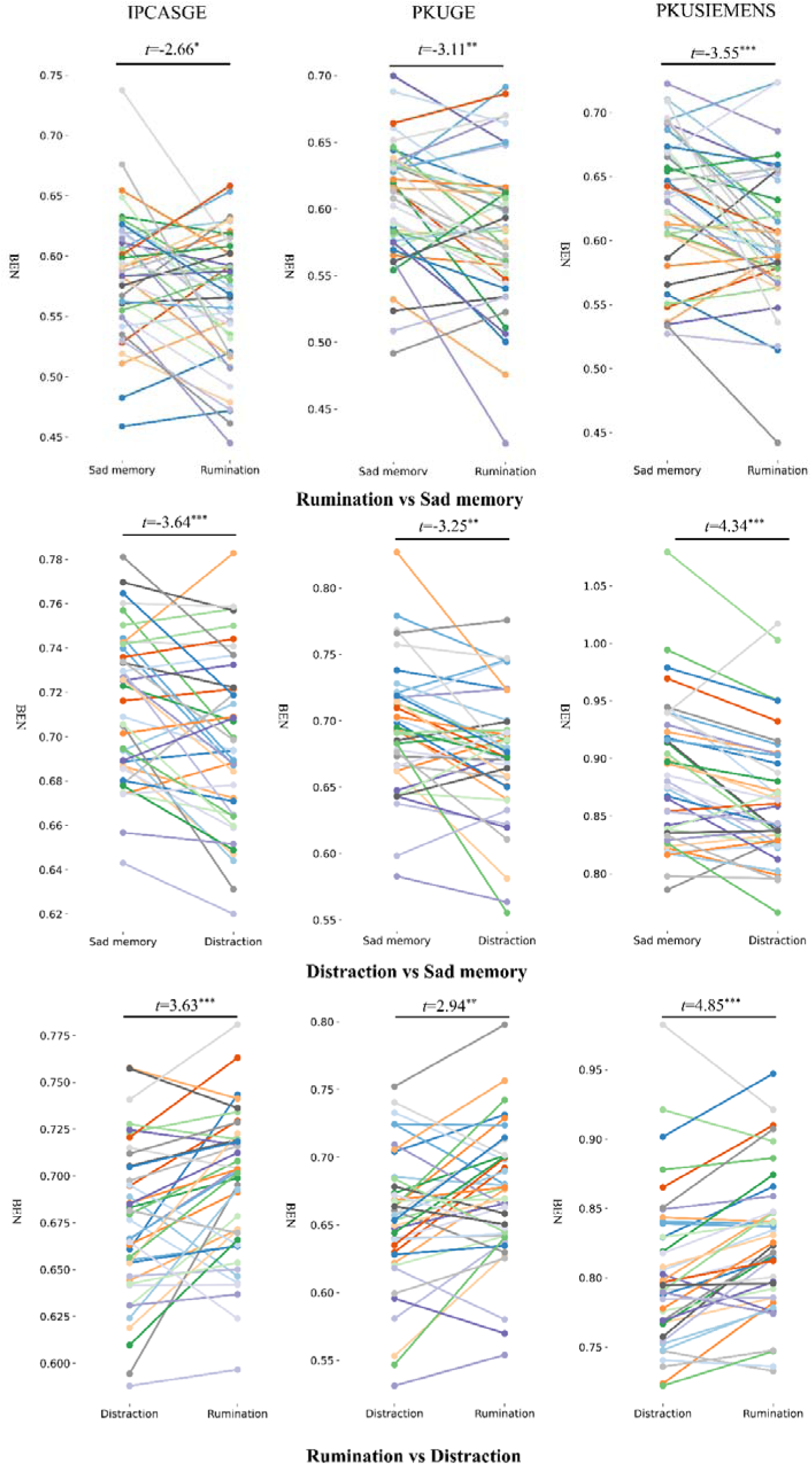
The mean BEN values were extracted from the overlapping brain regions for each subject and each scan. The x-axis represents different task states, while the y-axis represents the mean BEN values. The line colors distinguish between different subjects. T values from paired t-test, significance levels * *p* < 0.05, \*\**p* < 0.01, and \*\*\**p* < 0.001.

## 4. Discussion

In this study, we first observed that, compared to the resting state, task states showed increased BEN in the transmodal association cortices, including the DLPFC, TPJ, and PCC/PCu. In contrast, lower BEN was observed in the unimodal VC. Then lower BEN in VC was found during rumination compared to sad memory. Importantly, we observed decreased BEN during distraction compared to sad memory and increased BEN in the PCC/PCu during rumination compared to distraction. The PCC/PCu is a core hub of DMN, which is implicated in self-referential and internally focused cognitive processes (Harrison, Pujol et al. 2008, Sheline, Barch et al. 2009, Andrews-Hanna, Smallwood et al. 2014, Utevsky, Smith et al. 2014, Raichle 2015, Davey, Pujol et al. 2016, Liu, Nour et al. 2022, Menon 2023) and this finding is consistent with our hypothesis that rumination leads to heightened BEN in the DMN. The reproducibility of these results across multiple scanners and vendors underscores the robustness of the observed effects measured from BEN.

BEN in the VC was reduced during task states compared to the resting state, and lower BEN during rumination rather than distraction compared to sad memory. This pattern may reflect a shift in focus during tasks, where attention is directed toward task content, reducing the processing of external environmental information. In particular, rumination involves repetitive, internally focused thoughts on negative events, which likely decreases the demand for external information processing, resulting in lower BEN in the VC. Consistently, lower BEN in VC was expected during rumination than distraction, although this result was not replicated across all three scanners, we observed significantly lower BEN in the VC during rumination compared to distraction on two scanners (PKUGE and PKUSIEMENS) (Fig. 1D). This result is further supported by a study showing that self-reported rumination tendency is associated with reduced activation in the VC in non-depressed individuals (Piguet, Desseilles et al. 2014). In depression, abnormal functioning of the VC has been reported, and its activity is closely associated with both the onset of depression and the efficacy of antidepressant treatments (Furey, Drevets et al. 2013, Liu, Song et al. 2020, Wu, Lu et al. 2023) . A study has shown that individuals with MDD exhibit lower VC activity at baseline compared to healthy controls. Furthermore, lower VC activity during emotional processing tasks is associated with a better treatment response to scopolamine in MDD (Furey, Drevets et al. 2013) and our previous research on MDD found that greater BEN increase in the VC is associated with better disease improvement after 8 weeks of treatment (Liu, Song et al. 2020).

Increased BEN in the PCC/PCu was observed during rumination compared to the distraction. No significant change in BEN was found in the PCC/PCu during rumination when compared to the sad memory. In contrast, lower BEN was observed during distraction relative to the sad memory. These findings reflect that, during rumination, individuals remain deeply engaged in the repetitive processing of negative events, while during distraction, attention is redirected away from self-referential and internally focused cognitive processes. This result is consistent with other studies that have found a close association between the PCC/PCu and rumination (Cooney, Joormann et al. 2010, Berman, Peltier et al. 2011, Burkhouse, Jacobs et al. 2017, Chen, Chen et al. 2020, Philippi, Pessin et al. 2020, Zhou, Chen et al. 2020, Chen, Fan et al. 2024, Langenecker, Schreiner et al. 2024). Burkhouse et al (2017) found that all participants including remitted MDD and healthy controls recruited PCC during rumination, and increased activation in PCC during rumination was correlated with increased self-report rumination and symptoms of depression across all participants in adolescents (Burkhouse, Jacobs et al. 2017). Currently, using intracranial electroencephalogram (iEEG), Chen et al (2024) found that enhanced low-frequency power in the precuneus during the rumination condition compared to the control condition (Chen, Fan et al. 2024). Our recent research found increased BEN in the PCu in individuals with mild to moderate depression compared to healthy controls, which was subsequently suppressed following non-pharmacological treatment and another study also identified increased BEN in the right PCu in MDD relative to healthy controls, which was reduced after electroconvulsive therapy (Fan, Zhang et al. 2024).

In conclusion, this study consistently demonstrates decreased BEN in VC during rumination compared to sad memory and increased BEN in the PCC/PCu during rumination compared to distraction, suggesting that rumination is associated with a heightened focus on the internal world and a reduced processing of external information. This finding aligns with the concept that rumination involves repetitive, self-referential thinking, particularly focused on negative aspects of the self. The use of BEN provides a novel perspective for understanding the neural underpinnings of rumination and its role in depression and other related mental disorders.

## Acknowledgments

We thank Xiao Chen et al for releasing their dataset.

## Data and code availability

The original data is available from RMP Rumination fMRI Dataset (http://rfmri.org/RuminationfMRIData or http://doi.org/10.57760/sciencedb.o00115.00002).

The BEN maps and unthresholded statistical maps can be accessed from OSF (https://osf.io/r42yv/?view_only=3d82a0a04db04983957e0542c99a531a).

BENtbx is available at https://www.cfn.upenn.edu/zewang/BENtbx.php and https://github.com/zewangnew/BENtbx.

Nilearn is available at https://nilearn.github.io/dev/index.html.

## CRediT authorship contribution statement

Jue Lu: data analysis, visualization, manuscript drafting, Donghui Song: conceptualization, data analysis, visualization, manuscript review and editing. Da Chang: manuscript review and editing, Ze Wang: conceptualization, manuscript review and editing, supervision.

